## Supplemental Information for "Properties of a Multidimensional Landscape Model for Determining Cellular Network Thermodynamics"

**This PDF file includes:**

**Supplementary Text**

### Supplemental Information 1. Thermo-Fokker Planck (Thermo-FP) dynamics

Rationale for the use of Thermo-FP

In a large population of cells in which phenotypic characteristics such as gene expression product levels can be treated as continuous variables, the usefulness of a Thermo-FP description resides in its economy and generality. Knowledge of the N-component drift vector and the N x N diffusion matrix are sufficient to specify a unique probability density $W(\{x\},t)$. The drift vector represents the influence of the deterministic force (i.e. the gradient of the potential) and the diffusion matrix determines the rate and magnitude of stochastic fluctuations. Together the drift vector and diffusion matrix describe how the distribution changes in time in the Fokker-Planck (FP) equation. These factors are experimentally accessible from the observed steady state distribution and the measured dynamic correlations (diffusion coefficients) between variables. We have also shown that changes in the population distribution during relaxation allows estimation of the diffusion correlations, $D_{ij}$, directly from the Boltzmann H-function (1). The Boltzmann H-function relates the change in the population distribution during relaxation to the decrease in free energy and the resultant dissipative heat. The positive definite form of $-\frac{dH(t)}{dt}$ associated with FP dynamics involves binary products of gradients of a chemical potential, $\mu$, defined by

$$\mu = ln\frac{W(\{x\},t)}{W_{ss}(\{x\})}$$

This implies that Thermo-FP dynamics satisfies a variational principle, in that the steepest descent or most direct "path" from $W(\{x\},t)$ to $W_{ss}(\{x\})$ is always followed, and $-\frac{dH(t)}{dt}$ is as large as possible given the constraints specified by $D_{ij}(\{x\})$. The rate of degradation of free energy into heat is maximal throughout the relaxation process. In contrast to this dynamic behavior, the stationary state density $W_{ss}(\{x\})$ is one of minimal rate of dissipation and minimal rate of entropy production, so that homeostasis is guaranteed to be maximally efficient in a thermodynamic sense. The twin features of maximal rate of relaxation and maximum operating efficiency in the stationary state (steady state) make Thermo-FP dynamics an attractive option for characterizing cell population dynamics and thermodynamics.

Applying Thermo-FP dynamics to relaxation to the steady state

We define the relative free energy associated with a pair of probability densities, ($W_1$, $W_2$) as

$$H(t,\tau) = \int dx^N\, W_1(\{x\},t)\ln\left[\frac{W_1(\{x\},t)}{W_2(\{x\},t+\tau)}\right] \tag{S1}$$

where $t$ and $t + \tau$ are two time points and {x}=($x_1$,$x_2$,…$x_N$) is a set of N-dimensional coordinates. It should be emphasized that H is invariant with respect of coordinate representation (i.e. {x} → {x*}), which implies a high degree of flexibility in measurement protocols.

Now suppose ($W_1$, $W_2$) are solutions to the Thermo-FP equation (2)

$$\frac{\partial W(\{x\},t)}{\partial t} = \mathcal{L}_{FP}(\{x\},t)W(\{x\},t) \tag{S2}$$

Where

$$\mathcal{L}_{FP} = -\sum_{i=1}^{N} \frac{\partial}{\partial x_i} \mathcal{D}_i\,(\{x\},t) + \frac{1}{2}\sum_{i,j}^{N} \frac{\partial^2}{\partial x_i \partial x_j} D_{ij}(\{x\},t) \tag{S3}$$

Here the operator $\mathcal{L}_{FP}$ incorporates the drift vector $\mathcal{D}_i$ (the gradient of a chemical potential) together with the positive definite diffusion tensor (matrix) $D_{ij}$, which we assume to be independent of time. By differentiating H in Eq. S1 with respect to *t* while setting $W_2 = W_{ss}(\{x\})$ where $W_{ss}(\{x\})$ refers to the time-invariant stationary (steady) state, we arrive at

$$\frac{dH(t)}{dt} = -\frac{1}{2}\sum_{i,j}^{N} \int dx^N W(\{x\},t) \frac{\partial}{\partial x_i} \ln\left[\frac{W(\{x\},t)}{W_{ss}(\{x\})}\right] \cdot D_{i,j}(\{x\}) \cdot \frac{\partial}{\partial x_j} \ln\left[\frac{W(\{x\},t)}{W_{ss}(\{x\})}\right] \tag{S4}$$

The negative of Eq. S4, $-\frac{dH(t)}{dt}$, can be regarded as a positive definite quadratic form

$$-\frac{dH(t)}{dt} \geq 0 \tag{S5}$$

An important consequence of Eq. S4 is that it guarantees the stability of the steady state probability, $W_{ss}(\{x\})$, with respect to all transient perturbations (3). The stability of $W_{ss}$ is not restricted to small perturbations. A system evolving according to FP dynamics retains a perfect "memory" of its landscape, $-\ln W_{ss}(\{x\})$.

<u>Applying Thermo-FP dynamics to transitions of populations from one steady state to another</u>

In the current work we address a steady state, or stationary, distribution of phenotypic expression, through relaxation, involving deterministic drift and stochastic diffusion, to that steady state. We present an experimentally tractable theoretical approach using Thermo-FP dynamics to characterize the thermodynamics associated with correlated fluctuations between the network variables that maintain the steady state phenotype probability distribution. This approach can also be used to examine the role of network variables during a transition from one stationary state to another stationary state such as from a pluripotent to differentiated state. Population transitions can be defined by the Boltzmann H-function, where instead of $W_1(\{x\})$ and $W_2(\{x\})$ in Eq. S1 indicating distributions of the population at different points in time during the process of relaxation to the steady state, they represent different points in time during transition from one steady state to another.

Comparison of Thermo-FP with other approaches

The Thermo-FP approach that we present here can be contrasted with the optimal transport (OT) and population balance analysis (PBA) approaches to characterizing transitions (4, 5) by single cell (ss)RNA-seq analysis. All approaches describe the time evolution of a probability density of a population. FP dynamics is applicable to both population-based data in the form of coarse-grained time dependent probability densities, and to ensembles of single cell trajectories in the form of dynamic transition probability densities. Both our approach and PBA use FP dynamics which assumes that the evolution is dependent on both a deterministic drift vector and a stochastic diffusion component. Our construction, Thermo-FP, employs the H-function, which is equivalent to the Kullback-Leibler (KL) distance and corresponds to a relative free energy. This construction is a thermodynamic functional.

In our work stochastic diffusion is manifested in an N x N diffusion matrix consisting of diagonal (variance) and off-diagonal (covariance) elements of the dynamic fluctuations in network variables. This handling of diffusion preserves the connection with irreversible thermodynamics but comes at the considerable cost of having to measure fluctuations at every location in gene product expression space. PBA assumes a diffusion coefficient that is isotropic and invariant over gene expression space. In OT, the stochastic fluctuations of individual cell expression are not explicitly addressed. These assumptions make the OT and PBA approaches well-suited for the very large and sparsely populated data sets associated with time-dependent ssRNA-seq; the gene expression products of individual cells can be measured in many cells at single time points, but each cellular measurement is a terminal one and expression changes in individual cells cannot be followed over time. However, these approaches lose connection to the mechanistic insight that can be provided by thermodynamics.

### Supplemental Information 2. Experimental Considerations

The measurement of dynamic correlations of many cellular variables over time in individual cells can be achieved with time-resolved fluorescence microscopy of live cells. For example, quantitative live cell imaging can be used to monitor fluorescence intensity data from a cell line that has been genetically edited to express two fluorescence protein reporters for transcription activity of two different genes. The two fluorescent proteins could be different, such as mCherry and GFP. The dynamic data from each individual fluorescent protein could then be used to calculate a diagonal element (i=j) of the diffusion matrix, $D_{ij}(\{x\})$, and the temporal covariation of the two reporters could be used to calculate off-diagonal elements (i≠j) of $D_{ij}(\{x\})$ matrix. In this way a single cell line expressing two fluorescent protein reporters for gene expression can be used to construct a 2x2 diffusion matrix. To construct a higher dimensional diffusion matrix with cell lines expressing only two reporters at a time, all pairwise combinations of reporter are required such that an N-dimensional diffusion matrix requires $\frac{N\cdot[N+1]}{2}$ reporter cell lines. The capability to monitor the dynamics of more than two reporters simultaneously would greatly reduce the number of reporter cell lines required.

Estimating $D_{ij}(\{x\})$ from live cell imaging data has been performed previously in a single dimension (6) using 344 single cell trajectories. In a single dimension, there are no off-diagonal covariance terms and only a single fluorescence reporter is needed. By comparison, multidimensional landscapes have covariance terms and will require significantly more dynamic data to sample the larger dimensionality. Advanced automated live cell imaging methods can rapidly image large fields of view (7-9). These approaches can microscopically image live cell samples at rates >1 $cm^2$/min and are anticipated to provide datasets sufficient for estimating parameters in Equations 6-9, including the multidimensional diffusion matrix, $D_{ij}(\{x\})$. Shown below is an example computation estimating the number of live cell trajectories that could be obtained using rapid imaging approaches:

Average # of cells per area = 250 cells/$mm^2$
Imaging rate = 1 $cm^2$/min
Interval for timelapse imaging = 5 min
--------------------------------------------------------------------------------
Potential # of live cells imaged = 250 cells/$mm^2$ x 1 $cm^2$/min x 5 min = 125,000 single cell trajectories.

In the Sisan et. al.(6), the level of the fluorescence protein expression was observed to be cell cycle dependent. It is anticipated that the expression level of many genes/proteins will be cell cycle dependent (10). When estimating the elements of $D_{ij}(\{x\})$, the average cell cycle trend can be identified and normalized (6). The scale of data acquisition in order to achieve a large number of single cell trajectories and map a multidimensional landscape and corresponding diffusion matrix, $D_{ij}(\{x\})$, will also require computational approaches for handling and analyzing large image which are now available, for example described by Bascsy et. al. (11).

### Supplemental Information 3. Convergence of the homeostatic heat to an upper bound: A geometric interpretation.

Consider the inequality

$$\sum_{i=1}^{N} \lambda_i V_i^2 \leq C_{UB}$$

(S6)

where the $\lambda_i$ represent the positive eigenvalues of the diffusion tensor and $V_i$ are the gradients of the N-dimensional landscape. If the inequality expressed by Eq. S6 converges to an equality, then the invariance of $\dot{Q}_{ENERGY}$ with displacement from the stationary state becomes an indication that all relevant dissipative components of the landscape have been incorporated. The convergence of this sum to an upper bound, $C_{UB}$, or to any other limit is a subtle issue that we shall address, but not fully explore in this section.

For instance, the positive definite quadratic form of Eq. S6 can be regarded as defining the interior of an N-ellipsoid (12)

$$\sum_{i=1}^{N} \frac{V_i^2}{a_i^2} = 1$$

(S7)

for

$$a_i = \sqrt{\frac{C_{UB}}{\lambda_i}}$$

where $a_i$ is the length of the *i*-th semi-axis, and the N-dimensional volume of the ellipsoid (12) is given by

$$Volume_N = \frac{2}{N} \frac{\pi^{\frac{N}{2}}}{\Gamma\left(\frac{N}{2}\right)} (a_1 \cdot a_2 \ldots a_{N)}$$

(S8)

And where $\Gamma$ is Euler's gamma function. As N becomes large for a given set of $a_i$, we can employ Sterling's asymptotic approximation (13) to obtain

$$Volume_N \cong \frac{1}{\sqrt{N\pi}}\left(\frac{2\pi e}{N}\right)^{\frac{N}{2}} (a_1 \cdot a_2 \dots a_{N)}$$

(S9)

We conclude that $Volume_N$ vanishes rapidly and the ellipsoid shrinks to a point on the landscape as

$$Volume_N \sim \frac{1}{\sqrt{N}}\left(\frac{1}{N}\right)^{\frac{N}{2}}$$

(S10)

This very rapid shrinking of $Volume_N$ with increasing dimension is in striking contrast with that of an N-parallelepiped. The volume of a parallelepiped (enclosing an N-ellipsoid) whose sides contact the semi-axes is given by the parallelepiped

$$Volume_N = 2^N(a_1 \cdot a_2 \dots a_{N)}$$

(S11)

which *diverges* as N→ ∞. If one were to randomly insert points into the interior of this p-piped, virtually all would reside in the region exterior to the enclosed ellipsoid, even though 2N contacts exist between their surfaces.

Now consider the parallelepiped as a discretized version of a volume element located on the landscape. As the dimension, N, increases the ratio of the volume of the enclosed ellipsoid to that of the parallelepiped rapidly vanishes. Moreover, a randomly chosen point in the interior of the shrinking ellipsoid tends to reside arbitrarily close to its surface, so that the quadratic sum inequality introduced by Eq. 9 rapidly converges to an equality. An implicit assumption here is that the landscape is "rough" or "corrugated" over small length scales, so that gradients are never negligible.

To get a sense of how fast this large N convergence can occur, consider the case where all of the $a_i$ are the same $a$, so that we are dealing with an N-ball. Inscribe this N-ball into an N-cube so that the ball just touches the cube sides. Let the cube have sides length =1 so that its N-volume is 1, and the enclosed N-ball has a radius = ½. It turns out that the fraction of the total volume outside the N-ball but inside the N-cube equals 0.95 when N=6.

One might also consider the concentric N-balls having radii of say (0.9 $a$, $a$). How large must N be so that 4/5 of the volume lies in the annular shell of thickness 0.1a? The answer turns out to be N=8The volume of an N-ball having unit radius is 5.264 when N=5, and this is maximal for all N. This implies that, in this case, convergence to a single limit C is slowest for N=5 whereas the volume is 2.550 for N=10, 0.382 for N=15, and 0.258 for N=20.

It should also be noted that the summation in Eq. S6 can converge to a limit other than $C_{UB}$ if the following conditions are met. Suppose $\sum_{i=1}^{P} \lambda_i < L$ where $L$ is independent of P (the trace of the diffusion tensor is bounded). Then if $V_i^2 \geq V_{i+1}^2 > 0$ and $Limit\ V_i^2 = 0$, then $\sum_{i=1}^{\infty} \lambda_i V_i^2$ converges (13) (the total landscape "roughness" is bounded). Now this convergence can depend on location in the landscape, which implies that $\dot{Q}_{ENERGY}$ can depend explicitly on time and can vary with distance from the stationary state.

### Supplemental Information 4. The dissipative heat sustaining homeostasis of a network has a lower bound.

We show below that under certain conditions the probability distribution for the nonequilibrium steady state (NESS) can be very similar to that of the equilibrium distribution, and under those conditions the relative free energy between the nonequilibrium and the equilibrium steady states will be minimal.

The following is a proof that the phenotype probability distribution, which is a nonequilibrium stationary state (NESS) distribution, closely resembles the detailed balance (db) or equilibrium distribution under certain conditions.

Let $P(x)$ be the NESS probability distribution.

Let $Q(x)$ be the db probability distribution.

Let $g(x)$ represent the local rate of free energy dissipation on a landscape defined by $-ln\ P(x)$.

Consider the relative free energy, $D$, defined by

$$D(P \parallel Q) = \sum_x P(x) \ln[\frac{P(x)}{Q(x)}]$$

(S12)

Where x is a discrete microstate on a landscape.

Constrain $P(x)$ by $\sum_x P(x) = 1$, and $\sum_x P(x)g(x) = K$ where $g(x)$ represents the free energy dissipation rate at microstate x in a biochemical reaction network, and $K$ is the total dissipation rate for the entire landscape and equivalent to $\dot{Q}_{ENERGY}$; these constraints define $P^*(x)$:

$$P^*(x) = Q(x) \cdot \exp[\alpha g(x) + \alpha_0]$$

(S13)

where $\alpha$is a constant (independent of $x$) Lagrange multiplier and $\alpha_0$ is required for normalization of $P^*(x)$, and $\sum Q(x) = 1$. Moreover, $D(P^* \parallel Q)$ is minimized by $P^*(x)$ in a variational sense, i.e., $D(P^* \parallel Q)$ is as small as possible, given a specific rate, $K$, of free energy dissipation.

Note that $D(P \parallel Q) = \alpha \cdot K + \alpha_0$ for $(\lambda, K > 0)$ and the Lagrange multiplier, $\alpha$, can be determined from

$$\frac{dD\ (P^* \parallel Q)}{dK} = \alpha$$

(S14)

This minimization of $D(P \parallel Q)$ with constraint(s) on $P$is the flipside of entropy maximization with the same constraint(s) on $P$. (14)

Suppose that g(x) = K at every x, i.e., g(x) is uniform over the entire multivariable landscape. Then since $\sum_x Q(x) = 1$, we must have $\alpha \cdot K + \alpha_0 = 0$ or $K = -\alpha_0/\alpha$. Moreover, $D(P^* \parallel Q) = D(Q \parallel Q) = 0$ and the relative free energy sustaining homeostasis vanishes. Under these conditions, the network/landscape functions with energy/entropy compensation and the non-equilibrium steady state and detailed balance distributions are identical.

However, generally speaking, the homeostatic heat generation rate will not be uniform over the landscape, i.e. $K \geq -\alpha_0/\alpha$. As $K$ becomes larger, $P(x)$ and $Q(x)$ are increasingly dissimilar. This means that the heat generation rate is bounded from below, and there is a finite dissipative gap between the NESS and db distributions.

At this point, it should be noted that Speck and Seifert (15) have shown that the homeostatic dissipative heat (housekeeping heat) plays a fundamental role in generalizing the Jarzynski fluctuation theorem so as to deal with the free energy difference between any pair of NESS states rather than two db states. It should also be noted that the minimum dissipative gap width has been calculated in the context of two-state symmetric molecular machines by Schneider (16). The author concludes that under isothermal conditions, this minimum dissipation rate equals the ambient temperature multiplied by the entropy of a symmetric two-state system; i.e., $k_B T \ln 2 \approx 0.693 k_B T$. The relative free energy $dD\ (P^* \parallel Q)$ must decrease upon introducing any form of coarse-graining in $x$, and the ultimate coarse-graining protocol is bivariate or two-state. The existence of a minimal dissipative gap also implies the existence of a principle of least dissipation that is independent of specific dynamical assumptions such as Markov processes or additive noise. This principle must also be valid for deterministic dynamics with any distribution of initial conditions(17, 18). What is required is the existence of constraints, one for the normalization of a probability distribution and the other for specifying an average homeostatic heat production rate. The connection between Schneider's result , our conclusions, and the principle of least dissipation for general NESS systems is a topic for a separate in-depth study.
